# Evaluation of *Peregrinus maidis transformer-2* as a target for CRISPR-based control

**DOI:** 10.1101/2024.03.01.582949

**Authors:** Yu-Hui Wang, Dina Espinoza Rivera, William Klobasa, Marcé D. Lorenzen

## Abstract

The corn planthopper, *Peregrinus maidis*, is an economically important pest of corn and sorghum. Here we report the initial steps towards developing a CRISPR-based control method, precision guided sterile insect technique (pgSIT), for this hemipteran pest. Specifically, we evaluated the potential of *transformer-2* (*tra-2*) as a target for sterilizing insects. First, we identified *tra-2* transcripts within our *P. maidis* transcriptome database and performed RNA interference (RNAi) to confirm functional conservation. RNAi-mediated knockdown of *Pmtra-2* in nymphs transformed females into pseudomales with deformed ovipositors resembling male claspers. While males showed no overt difference in appearance, they were indeed sterile. Importantly, the results were similar to those observed in another planthopper, *Nilaparvata lugens*. We also used CRISPR/Cas9 genome editing to assess the impact of *tra-2* knockout in injectees. CRISPR-mediated knockout of *Pmtra-2* had lethal effects on embryos, and hence not many injectees reached adulthood. However, mosaic knockout of *Pmtra-2* did impact female and male fertility, which supports the use of *tra-2* as a target for pgSIT in this hemipteran species.

## INTRODUCTION

Hemipteran pests damage crops not only directly by feeding but indirectly by transmitting plant viruses. Many plant viruses are economically important, such as banana bunchy top virus, cucumber mosaic virus, maize streak virus and tomato yellow leaf curl virus [1]. There are few control methods for viral diseases. One such method is control of the virus vectors, which in most cases are hemipterans. Insecticides are the main way to control hemipterans, but it is still challenging due to their biology. Because hemipterans use their proboscis to feed on plant sap, most insecticides that cannot be absorbed and distributed systemically within the plant do not affect these insects. They can be killed by some contact insecticides. However, since these insects generally reside in hidden locations (nooks and crannies of the plant), contact insecticides become less effective. In addition to the limited choices of insecticides, the chemicals themselves have resulted in several serious environmental problems. Therefore, alternative methods for controlling hemipteran vectors are sorely needed.

Sterile insect technique (SIT) is an eco-friendly control measure that involves the release of sterilized insects to suppress the pest population. Generally, sterilized insects are sorted and just the males are released, which provides better suppression efficiency [2, 3]. Several pests have been successfully controlled using SIT including the New World screwworm, Mediterranean fruit fly, melon fly and pink bollworm [4–6]. As an alternative to irradiation, CRISPR/Cas9 genome editing was recently used to sterilize insects [7]. This new method features the precision of CRISPR/Cas9 genome editing, which results in fewer negative effects on sterilized insects, hence the name – precision guided SIT (pgSIT). Specifically, pgSIT requires two homozygous strains, one expressing Cas9 nuclease and the other expressing guide RNAs (gRNAs) that target genes for female viability and male fertility. Each strain is healthy and fertile on its own. However, when the strains are crossed all resulting progeny inherit both the Cas9 and gRNA genes. This enables CRISPR-based genome editing to occur, thereby killing female offspring and rendering males sterile. In addition to its precision, the CRISPR/Cas9 system is the most straightforward genome editing technology to date. It opens new avenues for genetic engineering in many insects including hemipteran pests, such as the brown planthopper, pea aphid and glassy-winged sharpshooter [8–10], thereby paving the way for the development of genetics-based control methods like pgSIT.

Zhuo *et al.* used RNA interference (RNAi) to knockdown *transformer-2* (*tra-2*), a sex determination gene, in the brown planthopper, *Nilaparvata lugens*. They discovered that injection of double-stranded RNA (dsRNA) targeting *Nltra-2* into nymphs produced infertile pseudomales and infertile males [11]. In other words, knockdown of *tra-2* sterilized all progeny and caused masculinization of females. These features could make *tra-2* an excellent target for pgSIT in hemipterans. Therefore, we evaluated the potential of *tra-2* as a target in a closely related species, the corn planthopper, *Peregrinus maidis*. *P. maidis* is a pest of corn and sorghum. It is also the vector of maize mosaic virus and maize stripe virus. The combination of *P. maidis* and viral infection can significantly reduce crop yield [12]. An advantage of using *P. maidis* is that methods for CRISPR-based genome editing have already been worked out [13]. Additionally, a Cas9-expressing strain already exists (Klobasa *et al.*, paper in preparation). The goals of this study were to: i) identify the *tra-2* ortholog in *P. maidis*; ii) confirm the functional conservation of *Pmtra-2* using RNAi; iii) evaluate the potential of *tra-2* as a target for pgSIT in hemipterans through CRISPR/Cas9 genome editing.

## MATERIALS AND METHODS

### Insect rearing

Corn planthoppers were reared on 5-week-old corn plants (cultivar Early Sunglow, Park Seed Company, Greenwood, SC) and maintained in 30 cm x 30 cm x 60 cm cages covered by nylon-mesh screen (BioQuip Products Inc., Compton, CA) in an insectary room set at 25±1 °C, ∼70% RH and 14:10 (light:dark) photoperiod.

### *Transformer-2* (*tra-2*) identification and phylogenetic analysis

To produce the most comprehensive set of transcripts possible, sex-specific transcriptome assemblies were generated by separately combining RNA-Seq data from *P. maidis* females (SRR22197767 and SRR22197769) or males (SRR22197768, SRR22197770) [14]). Reads were imported into Lasergene software (DNASTAR, Madison, WI) for quality trimming, and *de novo* assembly into contigs using default settings. The resulting contigs were annotated using OmicsBox software (BioBam, Valencia, Spain) with a BLASTx E value cutoff of 10^-5^. The annotated transcriptome assemblies were each exported as FASTA files and used to generate male- and female-specific BLAST databases (BlastStation, TM Software, Arcadia, CA). The deduced amino-acid sequence of *N. lugens tra-2* [11] was used as a query sequence to search the female or male transcriptome database respectively. The same two *P. maidis* transcripts were top hits in both searches and based on the E value and percent identity they were selected for further examination (sequences can be found in the supporting information). The transcripts or their deduced amino-acid sequences (translated at https://web.expasy.org/translate/) were used as BLASTx or BLASTp queries respectively at NCBI (https://blast.ncbi.nlm.nih.gov/Blast.cgi) to confirm identity.

The alignment of Pmtra-2 and other tra-2 proteins was generated using SnapGene (GSL Biotech, San Diego, CA) with the MUSCLE algorithm. The phylogenetic tree was built using MEGAX [15]. The tra-2 amino-acid sequences were aligned using the MUSCLE algorithm and tested with 56 amino-acid substitution models. Then, the tree was constructed using the maximum likelihood method with the best-fitting substitution model (LG+G+F) and partial deletion treatment of gaps or missing data and re-sampled using 1000 bootstrap replicates.

### Expression of *Pmtra-2* across developmental stages

Total RNA was isolated from a range of *P. maidis* life stages, from mid-stage embryo to adult, using the Qiagen RNeasy mini kit (Qiagen, Hilden, Germany) and treated with DNase I (Qiagen, Hilden, Germany) according to the manufacturer’s instructions. Then, RNA was reverse-transcribed into cDNA using the SuperScript III first-strand synthesis kit (Thermo Fisher Scientific Inc., Waltham, MA). The cDNA was used as template in semiquantitative PCR with *Pmtra-2-* or *Ribosomal protein L10-* (*RPL10*, reference gene) [16] specific primers (MyTaq DNA Polymerase, Meridian Bioscience, Cincinnati, OH). The primers used are shown in S2 Table.

### RNAi-mediated knockdown of *Pmtra-2*

Double-strand RNA (dsRNA) targeting the common region of the two *Pmtra-2* transcripts was synthesized. Nested PCR was used to add T7 promoter sequences to the ends of the target fragment. In the 1^st^-round of PCR, the target fragment was amplified using cDNA as template along with target-specific primers. The amplification product was sized checked via agarose gel electrophoresis and then isolated from a section of the gel by soaking it in sterile water overnight. The overnight solution was used as template for a 2^nd^-round of PCR. In this round, each target-specific primer was flanked by the T7 promoter sequence. The resulting PCR product was purified using the QIAquick PCR purification kit (QIAGEN, Hilden, Germany) and sequenced to confirm the identity (S3). Then, 1 μg was used as template for dsRNA synthesis using the MEGAscript T7 transcription kit (Thermo Fisher Scientific Inc., Waltham, MA). The dsRNA was purified using the Direct-zol RNA miniprep kit (Zymo Research, Irvine, CA), and the concentration was determined using a NanoDrop 1000 Spectrophotometer (Thermo Fisher Scientific Inc., Waltham, MA). Phenol red (final concentration at 20%) was mixed with the dsRNA solution for ease of injection.

We chose to inject 3^rd^-instar nymphs as a means to balance ease of injection with the likelihood of generating observable phenotypic changes. Nymphs were anesthetized on ice, placed in an injection arena (9 cm-diameter Petri dish containing a layer of 1% agarose) and injected with ∼60 ng of *Pmtra-2* or *EGFP* dsRNA (negative control) using a FemtoJet microinjector (Eppendorf, Hamburg, Germany). Injectees were maintained in a two-ounce plastic cup with corn leaves until they became adults, and the resulting phenotypic changes were recorded. To confirm knockdown, four nymphs were collected on the 5^th^ day post injection for use in real-time quantitative reverse transcription PCR (qRT-PCR). RNA extraction and cDNA synthesis were conducted as described above (section: Expression of *Pmtra-2* across developmental stages).

Then, the cDNA was used as template in qRT-PCR (Maxima SYBR Green/ROX qPCR Master Mix; Thermo Fisher Scientific Inc., Waltham, MA) with *Pmtra-2-* or *RPL10-* (reference gene) [16] specific primers (S2 Table). The qRT-PCR was performed on a BioRad CFX384 C1000 Touch Thermocycler (BioRad Hercules, CA) using the following program: 95°C for 10 minutes, 40 cycles of 95°C for 15 seconds, 57°C for 30 seconds, 72°C for 15 seconds and a final dissociation curve step to check for non-specific amplification. The data was exported as a xlsx file using BioRad CFX Manager Software (BioRad Hercules, CA), and analyzed using Microsoft Excel 2016 (Microsoft, Redmond, WA).

If not specified, all PCR amplifications associated with RNAi-mediated knockdown of *Pmtra-2* were carried out using MyTaq DNA Polymerase (Meridian Bioscience, Cincinnati, OH). The primers used are shown in S2 Table. Thirty to sixty nymphs were injected in each treatment, and the experiment was performed three times.

We did further experiments to test the fertility of male injectees because male sterility is the key to pgSIT. After the injectees reached adulthood, each 1- to 5-day-old male was crossed to three young virgin wild-type females on a 2-week-old corn seedling for ten days. Then, the adults were removed from the plant, and the number of nymphs emerging was recorded in the following two weeks. The experiment was performed three times.

### CRISPR-mediated knockout of *Pmtra-2*

Embryo collection and CRISPR/Cas9 genome editing were conducted as described in the previous study [13]. In brief, precellular embryos were obtained by placing ∼15 females and ∼5 males into an agarose-based egg-laying chamber and collecting embryos ∼16 hours later. Embryos were injected with gRNAs (Synthego Corporation, Menlo Park, CA) and Cas9 nuclease (TrueCutTM Cas9 Protein v2; Thermo Fisher Scientific Inc., Waltham, MA). The injected embryos were maintained in a humidity chamber for around a week. Then, the hatchlings were moved to a 5-week-old corn seedling where they were reared to adulthood before screening them as adults for potential CRISPR-induced somatic changes in sex-related phenotypes and used in mating assays. Each 1- to 5-day-old adult was crossed to three young virgin wild-type mates on a 2-week-old corn seedling for two weeks. Then, the adults were removed from the plant, and the number of offspring was recorded in the following two weeks. To detect CRISPR-based genome editing, genomic DNA of each injectee was extracted using the E.Z.N.A. Tissue DNA Kit (Omega Bio-tek, Norcross, GA), and the target region was amplified by two rounds of PCR. PCR was conducted as described earlier (section: RNAi-mediated knockdown of *Pmtra-2*) but using the proofreading PrimeSTAR HS DNA Polymerase (Takara Bio, Shiga, Japan) and Ex Taq DNA Polymerase (Takara Bio, Shiga, Japan) respectively in the 1^st^- and 2^nd^-round PCR. The primers used are listed in S2 Table. The 2^nd^-round PCR product was purified using the QIAquick PCR purification kit (QIAGEN, Hilden, Germany), sequenced and submitted for Synthego Inference of CRISPR Edits Analysis (https://ice.synthego.com/#/). ICE analysis provides an easy assessment of insertion-deletion (indel) mutations generated by CRISPR, outputting the indels and their associated frequencies (% contribution). Specifically, if two percent of indel is detected, it means that two percent of all sequence reads from a specific sample contains the indicated change. A heterogeneous mix of indels can be detected. To test our hypothesis that knockout of *Pmtra-2* is lethal for late-stage embryos, 17 well-developed but unhatched injected embryos were collected, and their genomic DNA was extracted for PCR amplification, sequencing, and ICE analysis as described above.

## RESULTS

### *Transformer-2* (*tra-2*) identification and phylogenetic analysis

Two putative *tra-2* transcripts were identified within our *P. maidis* transcriptome databases. Their deduced amino-acid sequences were nearly identical, so the one having the lowest E value and highest percent identity to Nltra-2 in the BLAST search was selected for downstream analyses. Transformer-2 belongs to the serine/arginine-rich-like (SR-like) protein family, which is involved in RNA splicing. These proteins have two distinct domains, arginine/serine-rich (RS) domain and RNA-recognition motif (RRM). The alignment (Fig 1A) shows that Tra-2 contains one highly conserved RRM flanked by RS domains, suggesting a conserved role in binding and splicing of RNA. The phylogenetic tree (Fig 1B) indicates a close evolutionary relationship between Pmtra-2 and Nltra-2 protein, which also suggests functional conservation of this gene in the two species.

**Fig 1.**
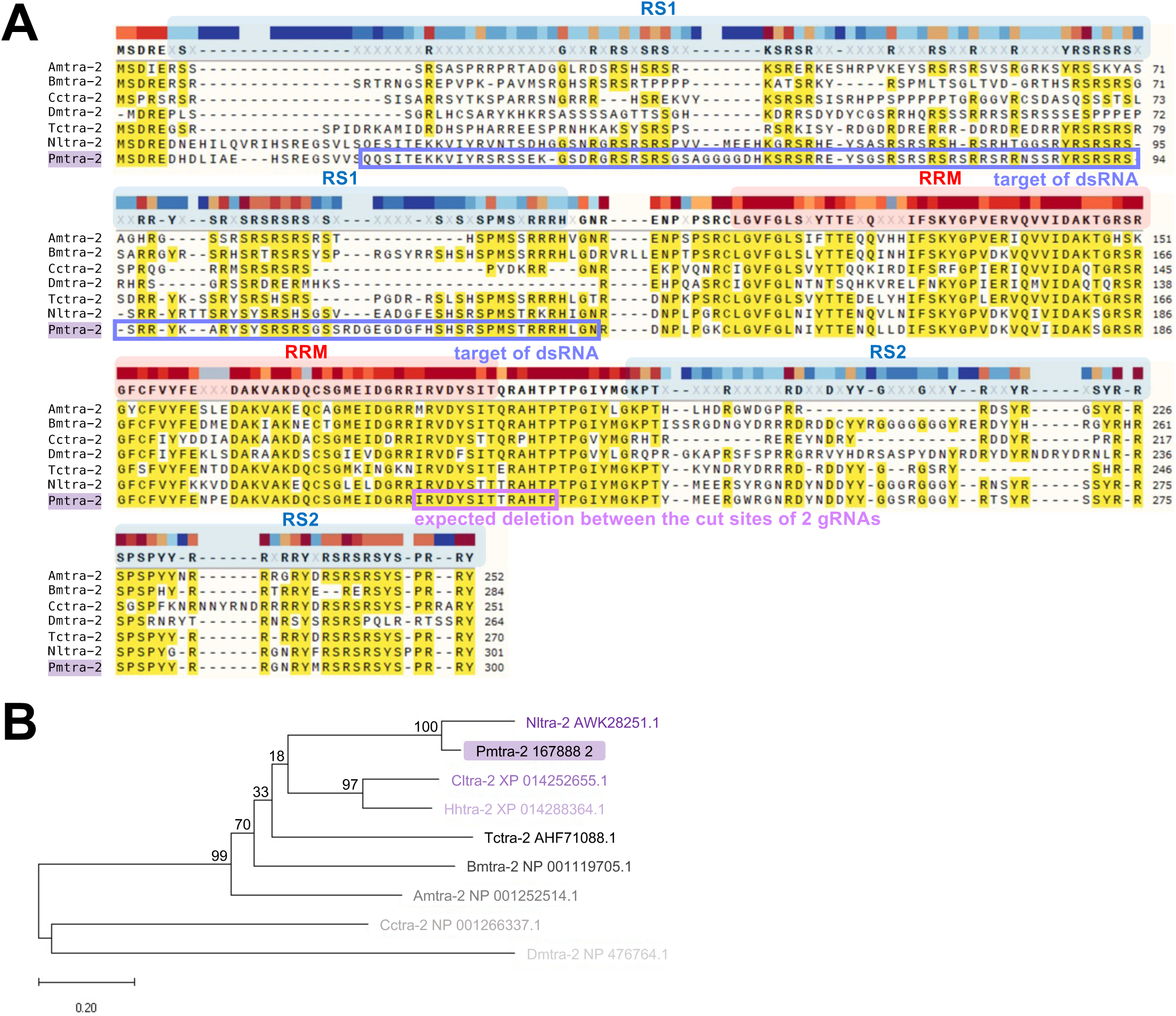
Alignment and phylogenetic tree of selected tra-2 proteins. (**A**) Alignment of the deduced amino-acid sequences of *tra-2* from *P. maidis* and other insects: *A. mellifera* (NP_001252514.1), *B. mori* (NP_001119705.1), *C. capitata* (NP_001266337.1), *Drosophila melanogaster* (NP_476764.1), *T. castaneum* (AHF71088.1) and *N. lugens* (AWK28251.1). The RS domains and RRM are marked. The target region of dsRNA and the expected deletion between the cut sites of two gRNAs are circled. (**B**) Phylogenetic tree based on the deduced amino-acid sequences of *tra-2* from *P. maidis*, *N. lugens* (AWK28251.1), *Cimex lectularius* (XP_014252655.1), *Halyomorpha halys* (XP_014288364.1), *T. castaneum* (AHF71088.1), *B. mori* (NP_001119705.1), *A. mellifera* (NP_001252514.1), *C. capitata* (NP_001266337.1) and *D. melanogaster* (NP_476764.1). The bootstrap values (%) are shown at the nodes. The scale bar represents an evolutionary distance of 0.2 amino-acid substitutions per site.

### RNAi-mediated knockdown of *Pmtra-2*

Since *Pmtra-2* is expressed across all developmental stages (Fig 2), we balanced the ease of injection with likelihood of phenotypic change [11] by injecting *Pmtra-2* dsRNA into 3^rd^- instar nymphs. Injectees were reared to adulthood, and external phenotypic changes in their reproductive state were assessed. Female injectees had deformed ovipositors that resembled male claspers, while males displayed no overt difference in their appearance (Fig 3A). It should be noted that injection of *EGFP* dsRNA did not impact *P. maidis* adult phenotypes. We further tested the fertility of male injectees by crossing each male to three young virgin wild-type females. As shown in Table 1, *Pmtra-2*-dsRNA-injected males produced almost no offspring, indicating that RNAi-mediated knockdown of *Pmtra-2* has a negative effect on male fertility. Importantly, the results were similar to those found in RNAi silencing of *tra-2* in *N. lugens* [11].

**Fig 2.**
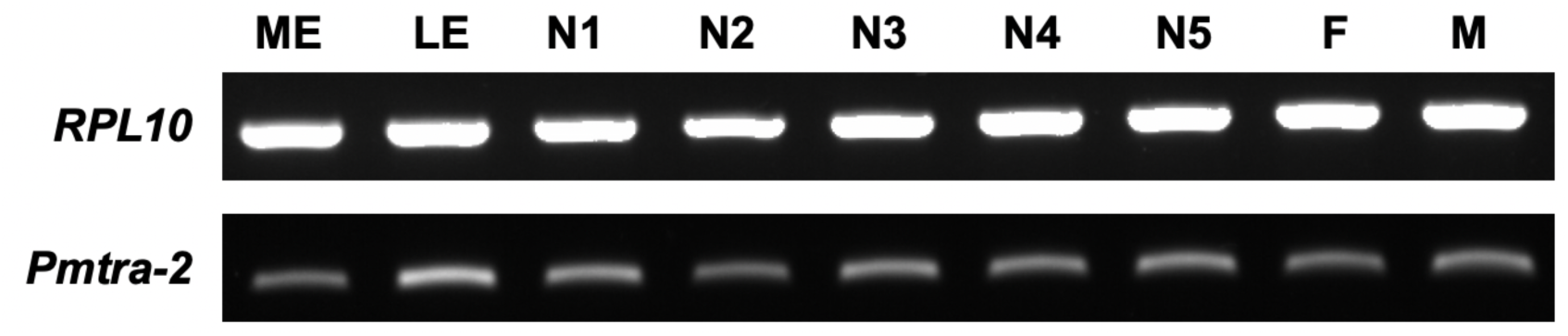
Expression of *Pmtra-2* across developmental stages. ME: middle-stage embryos, LE: late-stage embryos, N1: 1^st^-instar nymphs, N2: 2^nd^-instar nymphs, N3: 3^rd^-instar nymphs, N4: 4^th^-instar nymphs, N5: 5^th^-instar nymphs, F: adult females, M: adult males. Expression of *RPL10* was used as reference.

**Fig 3.**
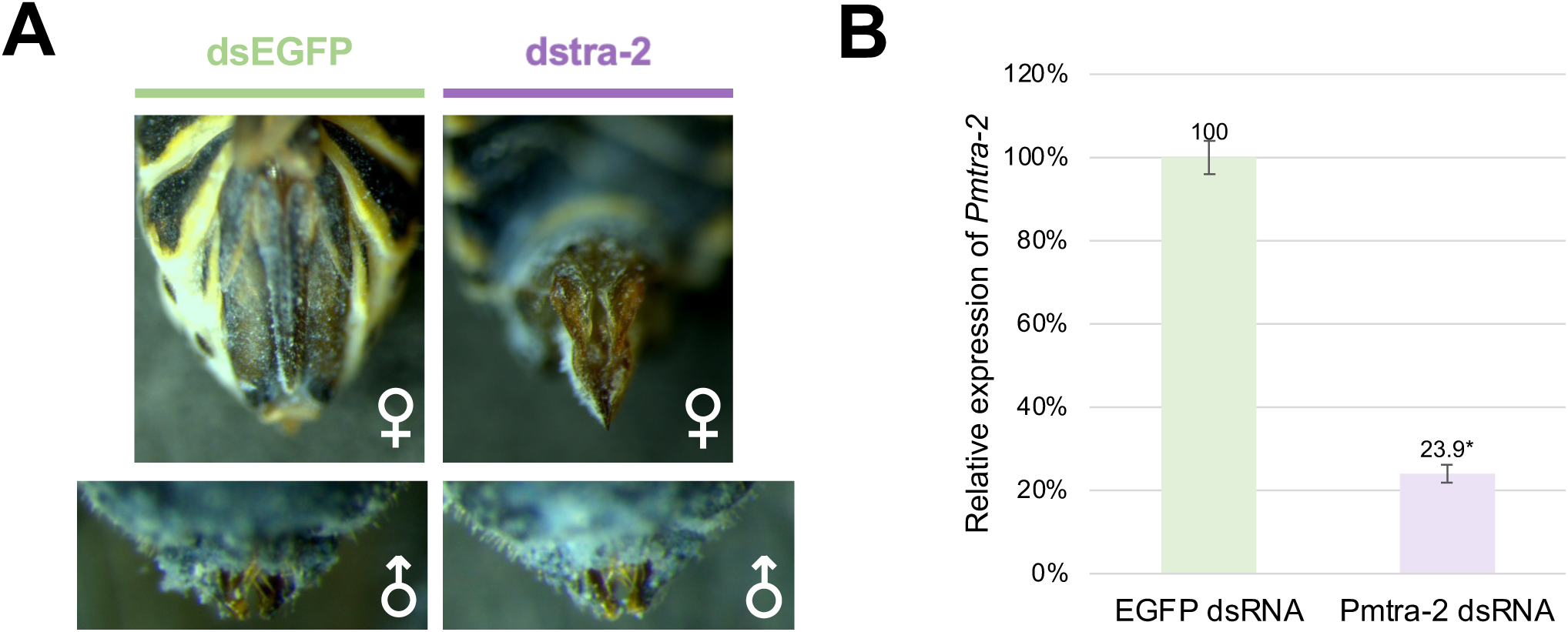
RNAi knockdown of *Pmtra-2* in nymphs. (**A**) Injection of *Pmtra-2* dsRNA transformed females into pseudomales with deformed ovipositors resembling male claspers, while there was no overt difference in male appearance. (**B**) Knockdown was confirmed by qRT-PCR. The *tra-2* transcript levels were normalized to the reference gene, *RPL10*. The asterisk above the bar indicates significant difference (P<0.05) as determined by Student’s t-test.

**Table 1.** The survival of dsRNA-injected nymphs and effect of knockdown on male fertility.

|  | Total number of injected nymphs | Nymphs reaching adulthood |  | Average number of offspring per male |
| --- | --- | --- | --- | --- |
|  |  | Males | Females |  |
| *EGFP* dsRNA | 34 | 9 | 7 | 77.4<br>(n=11) |
|  | 56 | 2 | 6 |  |
|  | 65 | 6 | 8 |  |
| *Pmtra-2* dsRNA | 48 | 6 | 0 | 0.3<br>(n=15) |
|  | 55 | 7 | 1 |  |
|  | 65 | 3 | 2 |  |

### CRISPR-mediated knockout of *Pmtra-2*

Screening for successful CRISPR/Cas9 editing events in the absence of a selectable marker can be like searching for a needle in a haystack. Therefore, given the sizeable amount of space, plants and personnel that would be needed to screen G_1_ progeny for individuals possessing a *tra-2* knockout allele, we opted to look at the impact of *tra-2* knockout in G_0_ injectees. Of the 925 embryos injected with Cas9 complexed with gRNAs targeting *Pmtra-2*, less than a third (259) showed signs of embryonic development. Of the 59 that hatched, only 30 survived to adulthood. Of these, only two adult females (09F and 11F) possessed deformed ovipositors (Fig 4). Since sterility is the key to pgSIT, we conducted mating assays to determine if fertility was impacted in any of the injectees. After giving injectees sufficient time to mate, we isolated their genomic DNA, amplified and sequenced the target region, and submitted the resulting sequence data for Synthego ICE analysis. The goal was to determine if changes in fertility were correlated with the levels of CRISPR-based genome editing. It should be noted that only injectees that survived until the end of the mating assay and who had at least two surviving mates are reported here (thus only 11). Interestingly, while the two females having deformed ovipositors (09F and 11F) failed to produce offspring (Table 2), ICE analysis detected little to no signs of genome editing. Specifically, no edited sequence was detected within the amplification products obtained from female 09F, and only two percent of the amplification products from female 11F possessed the 43-bp deletion (Fig 5). What was even more surprising than the extremely small percentage of genotypic change in the two females was the high levels of genotypic change in females having apparently normal ovipositors (1028F and 1031F; Table 2, Fig 5). Mosaic knockout of *Pmtra-2* did impact male fertility, though the quality of sequence obtained from some male injectees was too low for ICE analysis (Table 2).

**Fig 4.**
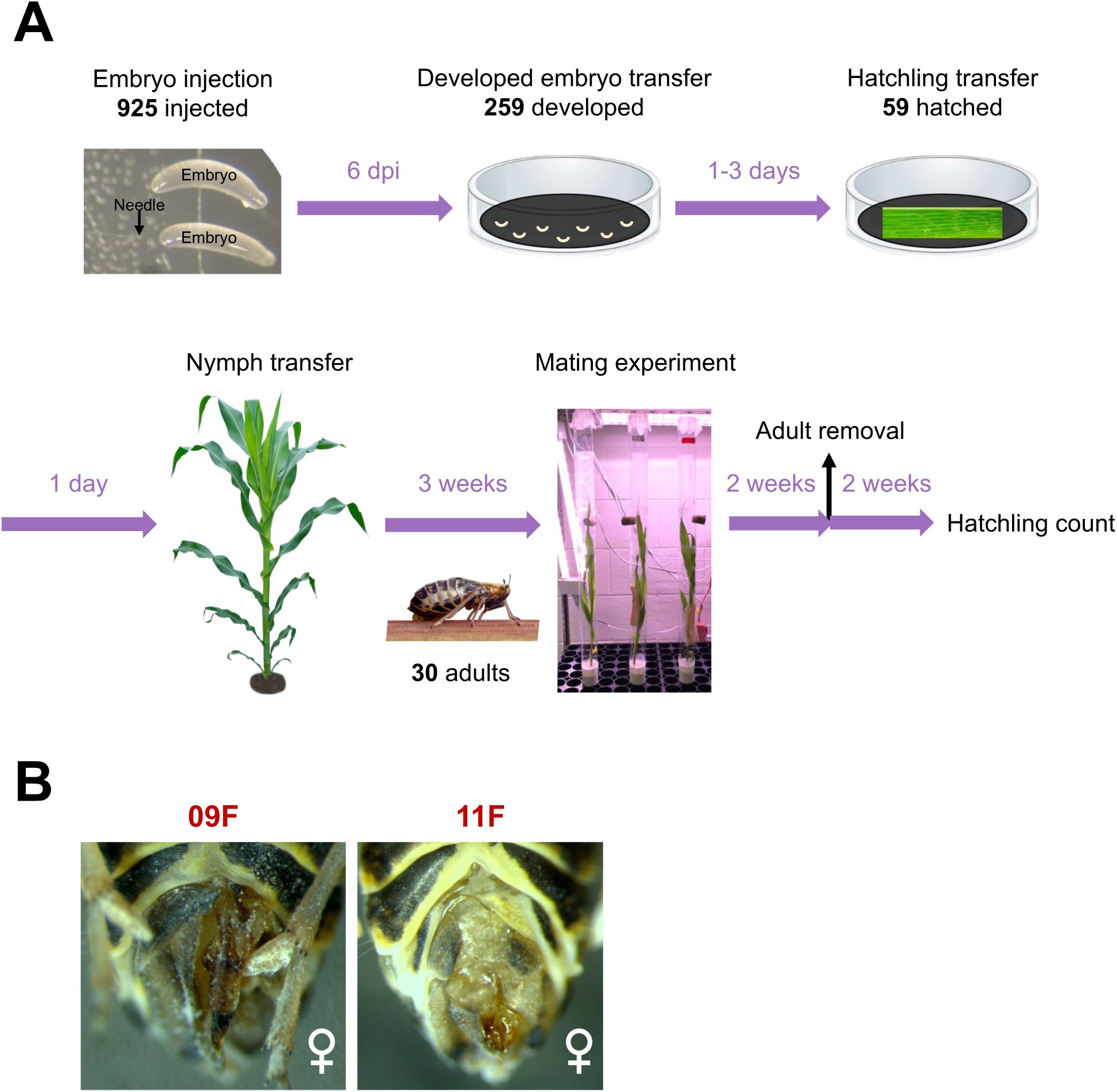
CRISPR-mediated knockout of *Pmtra-2*. CRISPR knockout was conducted to evaluate the potential of *tra-2* as a target for hemipteran pgSIT. (**A**) Schematic of workflow. (**B**) Two female injectees having deformed ovipositors.

**Fig 5.**
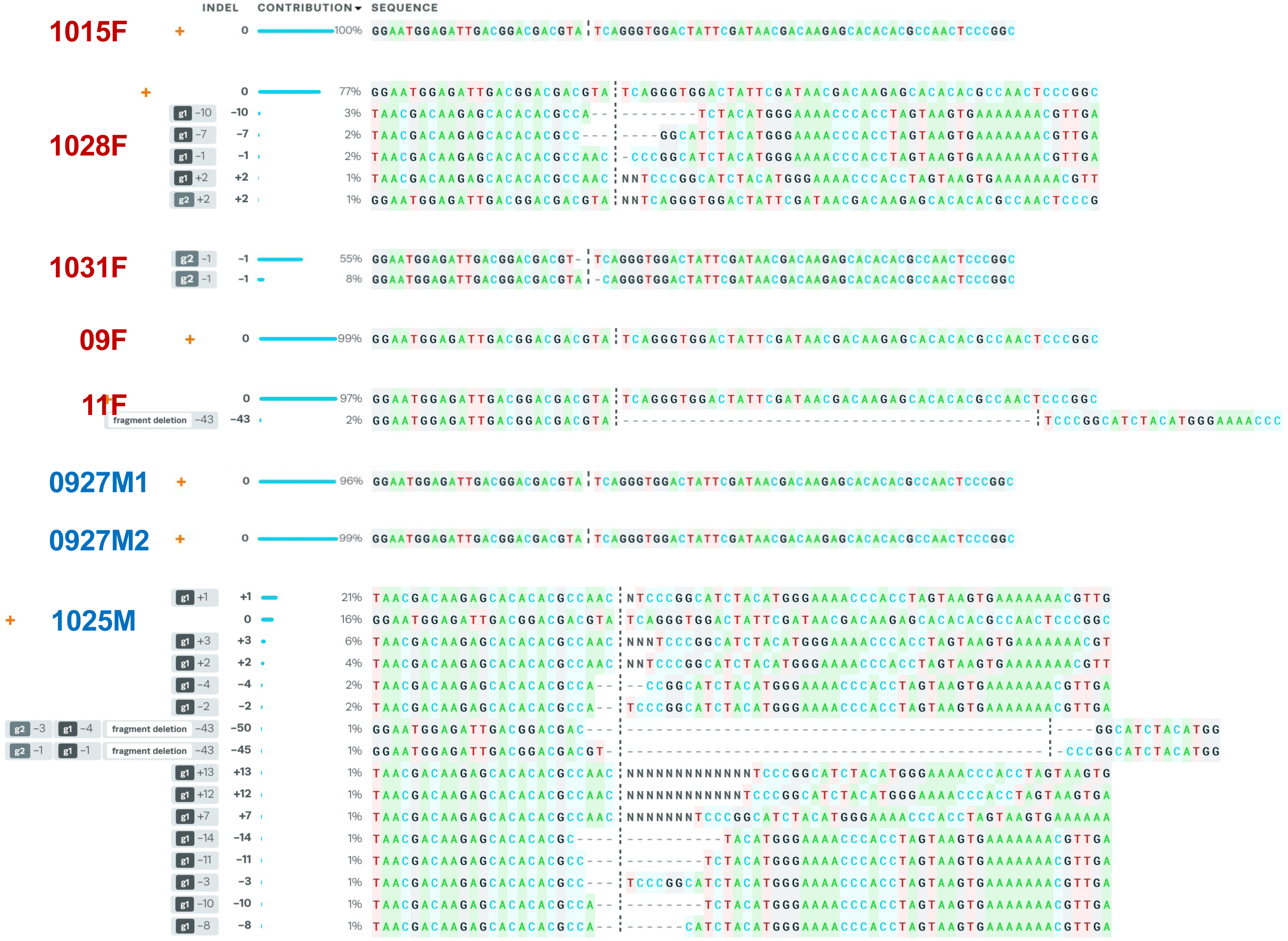
ICE analysis. The indels and their associated frequencies (% contribution) in injectees are displayed. The positions of indels (cut site of gRNA1/gRNA2) and the numbers of nucleotides inserted (+) or deleted (-) are indicated in the “Indel” column. The black vertical dotted lines represent the cut sites. The orange “+” symbols on the far left indicate unedited sequence.

**Table 2.**
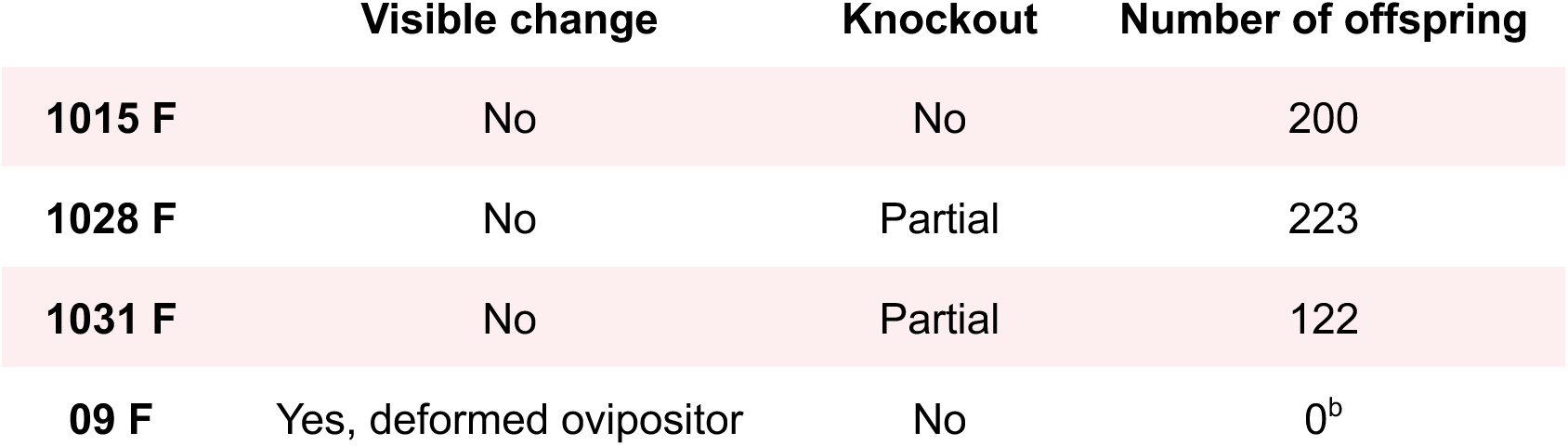
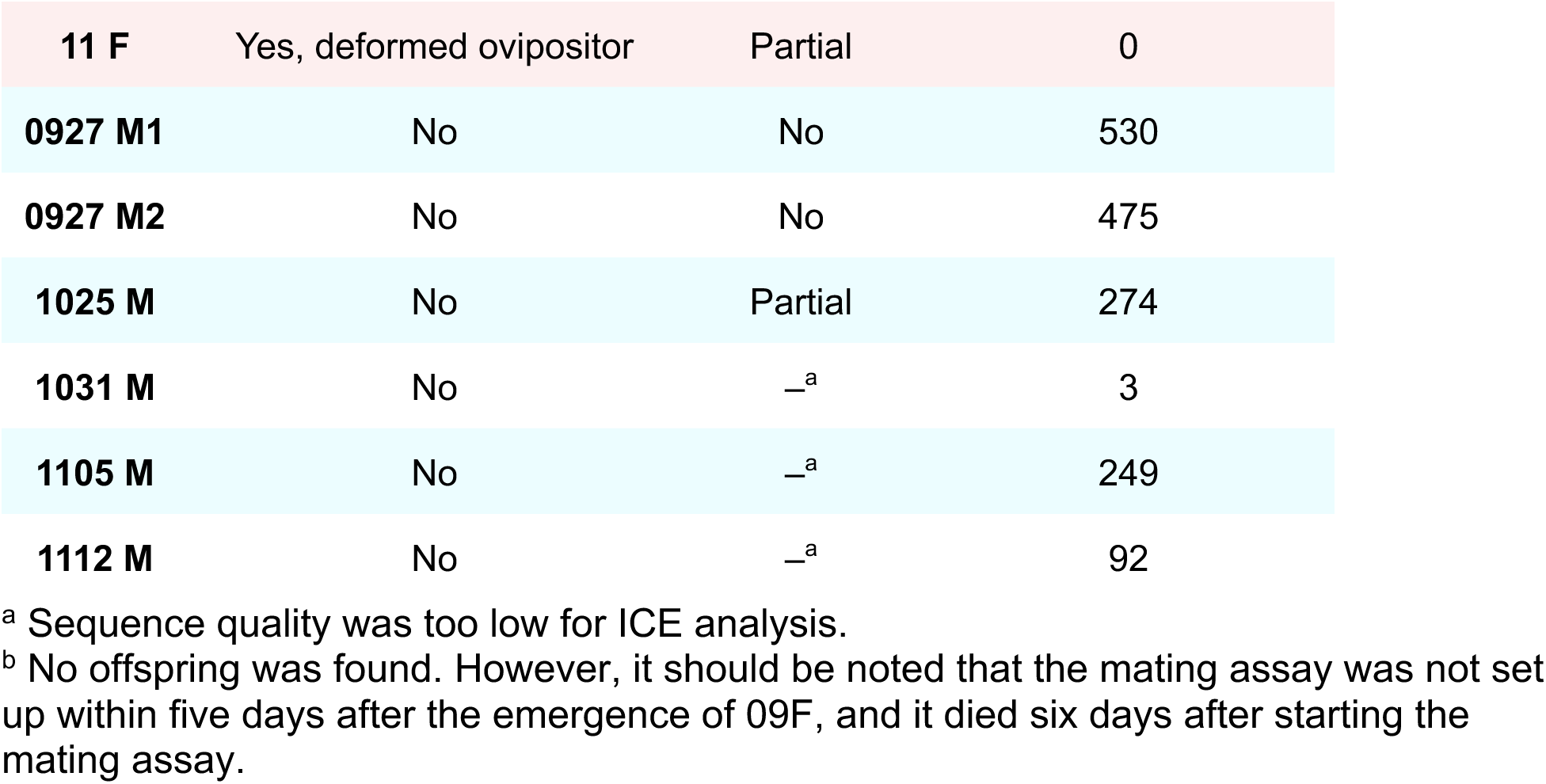
The result of mating assays.

Interestingly, we observed that many well-developed injected embryos (Fig 6A) died. Based on our experience, once embryos reach this stage of development, most hatch within two days. We believe that the low hatching rate (6.4%), especially for the well-developed embryos, was not due to injection because the hatching rate of embryos injected with buffer was 26.9% (of 119 injected embryos 32 hatched). Therefore, we hypothesized that high levels of *Pmtra-2* knockout is lethal in late-stage embryos. To test our hypothesis, we isolated genomic DNA from 17 well-developed but unhatched injected embryos, amplified and sequenced the target region, and submitted the sequence data for ICE analysis. The results indicate that our hypothesis is likely correct, since unhatched injectees had very high levels of edited sequences (Fig 6B).

**Fig 6.**
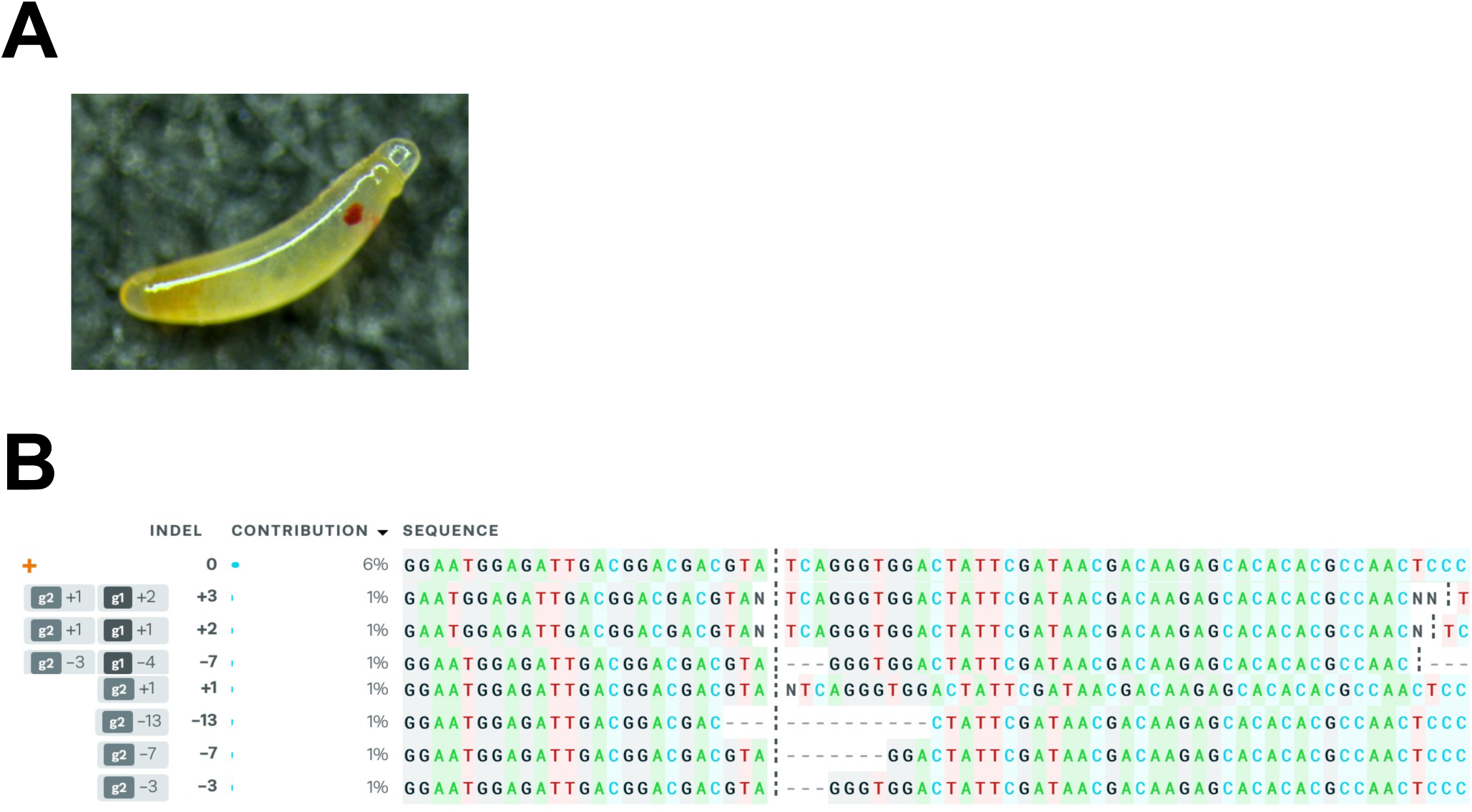
ICE analysis of 17 well-developed but unhatched injected embryos. (**A**) Developed but unhatched injectee. (**B**) ICE analysis output. The indels and their associated frequencies (% contribution) are displayed. The positions of indels (cut site of gRNA1/gRNA2) and the numbers of nucleotides inserted (+) or deleted (-) are indicated in the “Indel” column. The black vertical dotted lines represent the cut sites. The orange “+” symbols on the far left indicate unedited sequence.

## DISCUSSION

The number of *tra-2* transcripts varies between insects. For example, there are six known *tra-2* transcripts in *Bombyx mori* [17] and *Apis mellifera* [18], three in *Tribolium castaneum* [19], two in *N. lugens* [11], but only one in *Bactrocera dorsalis* [20] and *Diaphorina citri* [21]. Given the relatively close relationship between *P. maidis* and *N. lugens*, it is not surprising that the same number of *tra-2* transcripts was identified in both planthopper species. While the two *Nltra-2* transcripts only differ at the 5’ end, the *P. maidis* transcripts differ at the 5’ end as well as in the sequence connecting the RS1 and RRM regions.

Microinjecting early-stage nymphs is challenging but based on the data from Zhuo *et al.* [11] microinjecting *Nltra-2* dsRNA later in development resulted in weaker phenotypic changes. Therefore, we settled on delivering dsRNA to 3^rd^-instar nymphs, since individuals at this stage are large enough to handle, yet young enough to expect significant masculinization of females to occur. Indeed, our results were similar to those found in *N. lugens* [11]. Such results, masculinization of females due to *tra-2* knockdown, have been reported in a number of insects including *Musca domestica* [22], *Ceratitis capitata* [23] and *Anastrepha obliqua* [24]. There are differences though. In *N. lugens*, researchers also noticed a change in body size. Specifically, knockdown of *Nltra-2* resulted in smaller females and bigger males. However, this was not noted in other insects. We found that there were smaller females and bigger males, but these appeared to be individual differences, not the result of *Pmtra-2* knockdown.

To further evaluate the potential of *tra-2* as a target for pgSIT in hemipterans, we generated *tra-2* mutants using CRISPR/Cas9 genome editing. We used two gRNAs, one designed to target the conserved RS2 domain and the second targeting the linker sequence (Fig 1A), to maximize the chances of generating mutants by deleting the intervening sequence between the two cut sites. This also should have increased efficiency by compensating for the possibility that either gRNA might be ineffective. However, a deletion in this region could lead to a frameshift that would result in a new amino acid sequence at the 3’ end, and this could have a dominant negative effect on fitness. While female 09F and 11F had little to no signs of genome editing based on ICE analysis, they possessed deformed ovipositors. This suggests that in these individuals, *tra-2* may have only been knocked out in a small subset of tissues, thereby causing the level of edited sequences in female 09F to be too low for ICE analysis, but that small subset of tissues included the super important tissues involved in the development of the ovipositor. In comparison, female 1028F and 1031F had high levels of edited sequences but normal ovipositors likely because the genome-edited cells were not in the tissues that gave rise to their ovipositors. Taken together, the results suggest that where the genome-edited cells are located matters, which of course is not surprising.

CRISPR-mediated knockout of *Pmtra-2* impacted both male and female fertility, which is desirable for pgSIT, particularly male infertility. While mosaic knockout resulted in males producing fewer offspring, we were unable to determine the levels of genome editing in some injectees (1031M, 1105M and 1112M) due to sequence quality being too low for ICE analysis. It is possible that a highly heterogeneous mix of indels was created in these individuals, hence the low sequence quality. Another issue is that though a significant proportion of edited sequences was detected in male 1025M (only 16% wild type), it still produced offspring. While PCR amplification bias could be an explanation for the high levels of edited sequences, it is also possible that the genome-edited cells simply were not in its reproductive tissues. Either way, the lack of a complete *tra-2* knockout is problematic since we cannot be 100% sure that loss of Tra-2 function will result in sterile males, which is an absolute necessity for pgSIT.

We could not generate a complete *tra-2* knockout most likely due to lethality caused by loss of Tra-2 function during late-stage embryogenesis. Interestingly, similar finding was reported in *A. mellifera*. Namely, in honeybees injection of *tra-2* dsRNA into early-stage embryos resulted in death ∼70 hours post injection, which is also around the late embryonic stage [18, 25]. The results above suggest that *tra-2* plays a role in embryogenesis as well as in sex determination. This means that the timing of CRISPR-mediated *tra-2* knockout needs to be tightly controlled to avoid the lethal effects in the late embryonic stage if *tra-2* is to be used as a target for pgSIT.

Since our assay could not fully address questions regarding the outcome of CRISPR/Cas9-mediated knockout of *tra-2* due to only testing mosaic G_0_ injectees, future work will focus on knockout of *tra-2* and simultaneously knockin of a dominant marker gene such as DsRed. This would enable an easier screen since only the relatively rare G_1_ individuals expressing the fluorescent marker would need to be tested. Given limited resources (corn plants, space and personnel), it was not feasible to carry out a large-scale screen in the absence of a mechanism for detecting which G_1_ offspring should be tested via the rather time intensive crossing scheme. Importantly, the proposed screen would also allow us to test our hypothesis that knockout of *tra-2* is homozygous lethal.

As mentioned above, pgSIT requires two homozygous strains, one expressing Cas9 nuclease and the other expressing gRNAs targeting genes such as those involved in fertility. We currently have *P. maidis* strains that express Cas9 (Klobasa *et al.*, paper in preparation). However, since Cas9 is driven by a ubiquitin-like promoter, which drives constitutive expression, crossing these strains with a *Pmtra-2*-gRNA-expressing strain would likely kill all the offspring during the late embryonic stage. Therefore, to build a functional system, we plan to restrict Cas9 expression to later stages of development, or to limit expression to specific tissues, such as the gonads, by using age/tissue-specific promoters. We are also in the process of creating a *P. maidis* strain expressing *Pmtra-2* gRNAs. Given the desirable phenotypic changes observed in *Pmtra-2*-dsRNA-injected insects, an RNAi-based method of SIT could perhaps be used to control this hemipteran species [26].

## ACKNOWLEDGMENTS

We thank anonymous reviewers and the editor for the insightful comments and suggestions that improved our manuscript.

## AUTHOR CONTRIBUTIONS

Conceptualization, Y.-H.W. and M.D.L.; methodology, Y.-H.W., D.E.R. and M.D.L.; validation, Y.-H.W.; investigation, Y.-H.W., D.E.R. and W.K.; resources, M.D.L.; writing— original draft preparation, Y.-H.W.; writing—review and editing, Y.-H.W. and M.D.L.; visualization, Y.-H.W. and M.D.L.; supervision, Y.-H.W. and M.D.L.; project administration, Y.-H.W.; funding acquisition, M.D.L. All authors have read and agreed to the published version of the manuscript.

## REFERENCES

1. Rybicki EP. A top ten list for economically important plant viruses. Arch. Virol. 2015; 160: 17–20.

2. Rendón P, McInnis D, Lance D, Stewart J. Medfly (Diptera: Tephritidae) genetic sexing: large-scale field comparison of males-only and bisexual sterile fly releases in Guatemala. J. Econ. Entomol. 2004; 97: 1547–1553.

3. Vreysen MJ, Barclay HJ, Hendrichs J. Modeling of preferential mating in areawide control programs that integrate the release of strains of sterile males only or both sexes. Ann. Entomol. Soc. Am. 2006; 99: 607–616.

4. Calkins CO. Fruit Flies and the Sterile Insect Technique. 1st ed. Florida: CRC press; 1994.

5. Tabashnik BE, Liesner LR, Ellsworth PC, Unnithan GC, Fabrick JA, Naranjo SE, Li X, Dennehy TJ, Antilla L, Staten RT, Carrière Y. Transgenic cotton and sterile insect releases synergize eradication of pink bollworm a century after it invaded the United States. Proc. Natl. Acad. Sci. U.S.A. 2021; 118.

6. Hendrichs J, Vreysen MJB, Enkerlin WR, Cayol JP. Strategic options in using sterile insects for area-wide integrated pest management. In: Dyck VA, Hendrichs J, Robinson AS, editors. Sterile Insect Technique. 2nd ed. Florida: CRC press; 2021. p. 841–884.

7. Kandul NP, Liu J, Wu SL, Marshall JM, Akbari OS. Transforming insect population control with precision guided sterile males with demonstration in flies. Nat. Commun. 2019; 10: 1–12.

8. Xue WH, Xu N, Yuan XB, Chen HH, Zhang JL, Fu SJ, Zhang CX, Xu HJ. CRISPR/Cas9-mediated knockout of two eye pigmentation genes in the brown planthopper, *Nilaparvata lugens* (Hemiptera: Delphacidae). Insect Biochem. Mol. Biol. 2018; 93: 19–26.

9. Le Trionnaire G, Tanguy S, Hudaverdian S, Gléonnec F, Richard G, Cayrol B, Monsion B, Pichon E, Deshoux M, Webster C, Uzest M, Herpin A, Tagu D. An integrated protocol for targeted mutagenesis with CRISPR-Cas9 system in the pea aphid. Insect Biochem. Mol. Biol. 2019; 110: 34–44.

10. de Souza Pacheco I, Doss ALA, Vindiola BG, Brown DJ, Ettinger CL, Stajich JE, Redak RA, Walling LL, Atkinson PW. Efficient CRISPR/Cas9-mediated genome modification of the glassy-winged sharpshooter *Homalodisca vitripennis* (Germar). Sci. Rep. 2022; 12: 6428.

11. Zhuo JC, Lei C, Shi JK, Xu N, Xue WH, Zhang MQ, Ren ZW, Zhang HH, Zhang CX. Tra-2 mediates cross-talk between sex determination and wing polyphenism in female *Nilaparvata lugens*. Genetics 2017; 207: 1067–1078.

12. Autrey L. Maize mosaic virus and other maize virus diseases in the islands of the western Indian Ocean. In: Proceedings of the international maize virus diseases colloquium and workshop; 1983 August; Ohio.

13. Klobasa W, Chu FC, Huot O, Grubbs N, Rotenberg D, Whitfield AE, Lorenzen MD. Microinjection of corn planthopper, *Peregrinus maidis*, embryos for CRISPR/Cas9 genome editing. J. Vis. Exp. 2021; 169.

14. Wang YH, Klobasa W, Chu FC, Huot O, Whitfield A, Lorenzen M. Structural and functional insights into the ATP-binding cassette transporter family in the corn planthopper, *Peregrinus maidis*. Insect Mol. Biol. 2022; 32: 412–423.

15. Kumar S, Stecher G, Li M, Knyaz C, Tamura K. MEGA X: molecular evolutionary genetics analysis across computing platforms. Mol. Biol. Evol. 2018; 35: 1547–1549.

16. Yao J, Rotenberg D, Afsharifar A, Barandoc-Alviar K, Whitfield AE. Development of RNAi methods for *Peregrinus maidis*, the corn planthopper. PLoS One 2013; 8.

17. Niu BL, Meng ZQ, Tao YZ, Lu SL, Weng HB, He LH, Shen WF. Cloning and alternative splicing analysis of *Bombyx mori* transformer-2 gene using silkworm EST database. Acta Biochim. Biophys. Sin. 2005; 37: 728–736.

18. Nissen I, Müller M, Beye M. The Am-tra2 gene is an essential regulator of female splice regulation at two levels of the sex determination hierarchy of the honeybee. Genetics 2012; 192: 1015–1026.

19. Shukla JN, Palli SR. *Tribolium castaneum* Transformer-2 regulates sex determination and development in both males and females. Insect Biochem. Mol. Biol. 2013; 43: 1125–1132.

20. Liu G, Wu Q, Li J, Zhang G, Wan F. RNAi-mediated knock-down of transformer and transformer 2 to generate male-only progeny in the oriental fruit fly, *Bactrocera dorsalis* (Hendel). PLoS One 2015; 10.

21. Yu X, Killiny N. Effect of parental RNA interference of a transformer-2 homologue on female reproduction and offspring sex determination in Asian citrus psyllid. Physiol. Entomol. 2018; 43: 42–50.

22. Burghardt G, Hediger M, Siegenthaler C, Moser M, Dübendorfer A, Bopp D. The transformer2 gene in *Musca domestica* is required for selecting and maintaining the female pathway of development. Dev. Genes Evol. 2005; 215: 165–176.

23. Salvemini M, Robertson M, Aronson B, Atkinson P, Polito LC, Saccone G. *Ceratitis capitata* transformer-2 gene is required to establish and maintain the autoregulation of Cctra, the master gene for female sex determination. Int. J. Dev. Biol. 2003; 53: 109–120.

24. Ruiz MF, Milano A, Salvemini M, Eirín-López JM, Perondini AL, Selivon D, Sanchez L. The gene transformer of *Anastrepha* fruit flies (Diptera, Tephritidae) and its evolution in insects. PLoS One 2007; 2.

25. Cridge AG, Lovegrove MR, Skelly JG, Taylor SE, Petersen GEL, Cameron RC, Dearden PK. The honeybee as a model insect for developmental genetics. Genesis 2017; 55.

26. Darrington M, Dalmay T, Morrison NI, Chapman T. Implementing the sterile insect technique with RNA interference – a review. Entomol. Exp. Appl. 2017; 164: 155–175.

