## Supplementary for "Evaluation of *Peregrinus maidis transformer-2* as a target for CRISPR-based control"

### S1 Table. Primers and gRNAs used in this study.

| Use | Gene | Sequence (5' → 3') | Product length |
| --- | --- | --- | --- |
| **Primers**<br>dsRNA synthesis<br>the 1st-round PCR | *Pmtra-2* | F: tctcggttgtgtagatgttca<br>R: ttgtctcacagcagagcata | 420 bp |
| | *EGFP* | F: atggtgagcaagggcgaggagc<br>R: tttacttgtacagctcgtccatgc | 721 bp |
| **Primers**<br>dsRNA synthesis<br>the 2nd-round PCR | *Pmtra-2* | T7F: TAATACGACTCACTATAGGGtgtttccaagatggcgtc<br>T7R: TAATACGACTCACTATAGGGcagcagagcataacagagaa | 389 bp |
| | *EGFP* | T7F: TAATACGACTCACTATAGGGcacaagttcagcgtgtccg<br>T7R: TAATACGACTCACTATAGGGtgccgttcttctgcttgtc | 449 bp |
| **Primers**<br>qRT-PCR | *Pmtra-2* | F: acgatatcctcctcccct<br>R: gacgtatcagggtggact | 158 bp |
| | *RPL10* | F: cgccaacaagtacatggt<br>R: tccaaaggcacctctcat | 148 bp |
| **gRNAs** | *Pmtra-2* | gRNA1: AGAGCACACACGCCAACUCC<br>gRNA2: GAUUGACGGACGACGUAUCA | N.A. |
| **Primers**<br>knockout<br>confirmation | *Pmtra-2* | F: aggacaaccctctaccag<br>R: ttgcctgacagttcttgg | 492 bp |

**S1 File. *Pmtra-2* sequences.**

>Pmtra-2_167888_2
ccatgttttagaaaacagaacacacggaaccattttcttgtttgttgtatttgtgtattttctaatttggcgtcatcggtgttgctcata
aacttagattgcactcaaatttttagtaaggatatcttcaaatatgagtgatagagaagatcacgatcttattgctgagcacag
tcgagagggcagcgttgtctcacagcagagcataacagagaaaaaagtcatttataggtcgagatccagtgagaaggg
cagcgatcgtggcaggtcaaggtcacgttcgggaagcgccggtggtgggggagatcacaagagccgcagtagacgc
gagtattcgggctccagaagcagaagtagatcgcgcagtcgtagatcgcgtcgcaacagcagtcgctatcgctcgaggt
cgcgttcgtcgcgtcgctacaaggcacgctactcgtacagtcgctcgcgatctggatcgtcgcgcgatggagagggcgat
ggcttccattcgcactcgcggagtccgatgtcgaccaggcgacgccatcttggaaacagggacaaccctctaccaggca
agtgtctgggtgtgtttggtttgaacatctacacaaccgagaaccaattgttggacattttctcgaagtacggccccgttgaca
aggtgcaggtgattattgacgccaaatcgggccggtcgcgtggcttttgctttgtctatttcgagaatccggaagacgccaa
agtggcgaaagaccagtgctcgggaatggagattgacggacgacgtatcagggtggactattcgataacgacaagagc
acacacgccaactcccggcatctacatgggaaaacccacctacatggaagaacgcggatggaggggcaatagggatt
ataatgacgattactatggtggaagtaggggaggaggatatcgtaccagttacaggagcagttaccgacgatcaccgtcc
ccctactataggcggggcaatcgctacatgagatcaaggtcgcgctcatactcgccacgtcgttactaaactgtgatgggtt
ttcatgattgacgaatcagatgtcagacattggcttcctaacgtcgagatcctaacagctgactggaaagaaggctgactcc
cgttcagc

>Pmtra-2_167888_1
acagaacacacggaaccattttcttgtttgttgtatttgtgtattttctaatttggcgtcatcggtgttgctcataaacttagattgca
ctcaaatttttagtaaggatatcttcaaatatgagtgatagagaagatcacgatcttattgctgagcacagtcgagagggca
gcgttgtctcacagcagagcataacagagaaaaaagtcatttataggtcgagatccagtgagaagggcagcgatcgtgg
caggtcaaggtcacgttcgggaagcgccggtggtgggggagatcacaagagccgcagtagacgcgagtattcgggct
ccagaagcagaagtagatcgcgcagtcgtagatcgcgtcgcaacagcagtcgctatcgctcgaggtcgcgttcgtcgcg
tcgctacaaggcacgctactcgtacagtcgctcgcgatctggatcgtcgcgcgacggagagggcgatggcttccattcgc
actcgcggagtccgatgtcgaccaggcgacgccatcttggaaacagggtttccggctaccatggcgagagttatcaagat
aatcaacagtcaaaagacaaccctctaccaggcaagtgtctgggtgtgtttggtttgaacatctacacaaccgagaacca
attgttggacattttctcgaagtacggccccgttgacaaggtgcaggtgattattgacgccaaatcgggccggtcgcgtggct
tttgctttgtctatttcgagaatccggaagacgccaaagtggcgaaagaccagtgctcaggaatggagattgacggacga
cgtatcagggtggactattcgataacgacaagagcacacacgccaactcccggcatctacatgggaaaacccacctac
atggaagaacgcggatggaggggcaatagggattataatgacgattactatggtggaagtaggggaggaggatatcgt
accagttacaggagcagttaccgacgatcaccgtcccccctactagtaggcgggacggtgatcgtcggtaactg

The expected deletion between the cut sites of two gRNAs is in red.

**S2 File. Sequencing result of the 2nd-round PCR product.**

aaaagtcatttataggtcgagatccagtgagaagggcagcgatcgtggcaggtcaaggtcacgttcgggaagcgccggt
ggtgggggagatcacaagagccgcagtagacgcgagtattcgggctccagaagcagaagtagatcgcgcagtcgtag
atcgcgtcgcaacagcagtcgctatcgctcgaggtcgcgttcgtcgcgtcgctacaaggcacgctactcgtacagtcgctc
gcgatctggatcgtcgcgcgatggagagggcgatggcttccattcgcactcgcggagtccgatgtcgaccaggcgacgc
catcttggaaaca
